# Regeneration of the zebrafish retinal pigment epithelium after widespread genetic ablation

**DOI:** 10.1101/372151

**Authors:** Nicholas J. Hanovice, Lyndsay L. Leach, Kayleigh Slater, Ana E. Gabriel, Dwight Romanovicz, Enhua Shao, Ross Collery, Edward A. Burton, Kira Lathrop, Brian A. Link, Jeffrey M. Gross

**Author notes:** **Corresponding author:** Jeffrey M. Gross, Department of Ophthalmology, Louis J. Fox Center for Vision Restoration, University of Pittsburgh School of Medicine, Pittsburgh PA, 15213, USA.

## Abstract

The retinal pigment epithelium (RPE) is a specialized monolayer of pigmented cells within the eye that is critical for maintaining visual system function. Diseases affecting the RPE have dire consequences for vision, and the most prevalent of these is atrophic (dry) age-related macular degeneration (AMD), which is thought to result from RPE dysfunction and degeneration. An intriguing possibility for treating RPE degenerative diseases like atrophic AMD is the stimulation of endogenous RPE regeneration; however, very little is known about the mechanisms driving successful RPE regeneration *in vivo*. Here, we developed a zebrafish transgenic model (*rpe65a*:nfsB-GFP) that enabled ablation of large swathes of mature RPE. RPE ablation resulted in rapid RPE degeneration, as well as degeneration of Bruch’s membrane and underlying photoreceptors. Using this model, we demonstrate for the first time that larval and adult zebrafish are capable of regenerating a functional RPE monolayer after RPE ablation. Regenerated RPE cells first appear at the periphery of the RPE, and regeneration proceeds in a peripheral-to-central fashion. RPE ablation elicits a robust proliferative response in the remaining RPE. Subsequently, proliferative cells move into the injury site and differentiate into RPE. BrdU pulse-chase analyses demonstrate that the regenerated RPE is likely derived from remaining peripheral RPE cells. Pharmacological inhibition of Wnt signaling significantly reduces cell proliferation in the RPE and delays overall RPE recovery. These data demonstrate that the zebrafish RPE possesses a robust capacity for regeneration and highlight a potential mechanism through which endogenous RPE regenerate *in vivo*.

**SIGNIFICANCE STATEMENT:** Diseases resulting in RPE degeneration are among the leading causes of blindness worldwide, and no therapy exists that can replace RPE or restore lost vision. One intriguing possibility is the development of therapies focused on stimulating endogenous RPE regeneration. For this to be possible, we must first gain a deeper understanding of the mechanisms underlying RPE regeneration. Here, we ablate mature RPE in zebrafish and demonstrate that zebrafish regenerate RPE after widespread injury. Injury-adjacent RPE proliferate and regenerate RPE, suggesting that they are the source of regenerated tissue. Finally, we demonstrate that Wnt signaling is required for RPE regeneration. These findings establish an *in vivo* model through which the molecular and cellular underpinnings of RPE regeneration can be further characterized.

## INTRODUCTION

The RPE is a polarized monolayer of pigment-containing cells that separates the retina from the choroid, and performs many critical functions for vision. Microvilli extend from the apical RPE surface and interdigitate with photoreceptor outer segments (POS), enabling the RPE to support photoreceptor health (1). The basal surface of the RPE abuts and helps to form Bruch’s membrane (BM), which, along with tight junctions between RPE cells, creates the blood-retina barrier and facilitates nutrient and ion transport between the retina and choriocapillaris (2, 3). Additionally, RPE pigment prevents light scatter by absorbing stray photons. Due to its importance in maintaining retinal function, diseases affecting the RPE have dire consequences for vision. Age-related macular degeneration (AMD) is one such disease, and is the third leading cause of blindness in the world (4, 5). AMD is commonly divided into two types: atrophic (dry) and exudative (wet). In the early stages of atrophic AMD, RPE cells in the parafovea become dysfunctional and progressively degenerate, and this is thought to result in death of parafoveal rods (6–8). Progressively, RPE dysfunction and degeneration spreads to the fovea, resulting in loss of cone photoreceptors, and ultimately, loss of high-acuity vision (9, 10). Exudative AMD occurs in a subset of atrophic AMD cases when choroidal vasculature invades the retina (10, 11).

Transplantation of stem cell-derived RPE has emerged as a possibility for treating AMD (12, 13), and clinical trials are currently underway (e.g. (14–18)). An unexplored complementary approach is the development of therapies that stimulate endogenous RPE regeneration. In mammals, RPE regeneration is limited and dependent upon the size of the injury (19); small lesions can be repaired by the expansion of adjacent RPE cells, (20, 21), but existing RPE cells are unable to repair large lesions (19, 22–25). In some injury paradigms, RPE cells proliferate but do not regenerate a morphologically normal monolayer (e.g. (21, 26, 27)). Indeed, RPE cells often overproliferate after injury, such as during proliferative vitreoretinopathy (PVR), where proliferative RPE cells invade the subretinal space and lead to blindness (28, 29). Recently, a subpopulation of quiescent human RPE stem cells was identified that can be induced to proliferate *in vitro* and differentiate into RPE or mesenchymal cell types (25, 30), suggesting that the human RPE contains a population of cells that could be induced to regenerate.

Little is known about the process by which RPE cells respond to elicit a regenerative, rather than pathological, response. Indeed, no studies have demonstrated regeneration of a functional RPE monolayer following severe damage in any system. The development of such a model is a critical first step to acquiring a deeper understanding of the molecular mechanisms underlying RPE regeneration. Zebrafish offer distinct advantages for this purpose: the development, structure and function of the zebrafish eye is similar to human, including a cone-rich retina; they are amenable to genetic manipulation and imaging, and they can regenerate neural tissues (e.g. (31–33)). However, it is unknown whether the zebrafish RPE is capable of regeneration. Here, we demonstrate that the zebrafish RPE possesses a robust capacity for regeneration and identify cellular and molecular mechanisms through which endogenous RPE regenerate *in vivo*.

## RESULTS

### RPE ablation results in AMD-like phenotypes

To develop an RPE injury model, we utilized a transgenic line in which an *rpe65a* promoter element drives expression of the nfsB-EGFP fusion protein in mature RPE (34) (*rpe65a*:nfsB-GFP; SI Appendix Fig S1A,B). nfsB is an *E. coli* nitroreductase that converts the prodrug metronidazole (MTZ) into a potent DNA crosslinking agent, leading to apoptosis in expressing cells (35, 36). *rpe65a*:nfsB-GFP embryos were treated with phenylthiourea (PTU) (37) to suppress melanin synthesis. To ablate the RPE, 5dpf larvae were removed from PTU and exposed to 10mM MTZ for 24 hours. After treatment, GFP^+^ cells degenerate (SI Appendix Fig S1E,F), nuclei in the outer nuclear layer (ONL) adjacent to ablated RPE become disorganized and POS morphology is disrupted (SI Appendix Fig S1E’,F’). Degeneration of GFP^+^ cells was accompanied by the absence of pigmentation after removal of PTU (SI Appendix Fig S1C compared to Fig S1F; Fig S2, p<0.0001).

In atrophic AMD, RPE dysfunction and degeneration leads to degeneration of underlying photoreceptors (PRs) (9, 10). To characterize the temporal dynamics of RPE and PR degeneration following MTZ treatment, sections were taken from larvae at 3, 6, 12, 18, 24 and 48 hours post injury (hpi) and stained for TUNEL (Fig 1, SI Appendix S3). While the ONL appears normal at 3hpi, TUNEL^+^ nuclei appear in the RPE (SI Appendix Fig S3A,B) and RPE apical microvilli began to degenerate. By 6hpi, RPE apical microvilli were absent (SI Appendix Fig S3C,D) and nuclear organization in the ONL began to deteriorate. At 12hpi, apoptosis significantly increased in the RPE (Fig 1E, p=0.016) and ONL nuclei became delaminated. By 18hpi, apoptosis in the ONL increased significantly (Fig 1F, p<0.0001) and GFP accumulated in blebs, a process which left regions of RPE devoid of GFP signal (Fig SI Appendix S3G,H). RPE apoptosis peaked at 24hpi (Fig 1E, p<0.0001). Apoptosis, while remaining significantly elevated when compared to controls, began to decrease in both layers by 48hpi (Fig 1E,F, p<0.0001 in RPE, p=0.0301 in ONL). Morphologically normal GFP^+^ RPE cells were absent from the RPE by 24hpi, and by 48hpi, all remaining GFP signal was contained in irregular GFP^+^ blebs, likely consisting of RPE cell debris. ONL nuclear lamination remained severely disrupted (SI Appendix Fig S3I-L). Non-transgenic siblings treated with MTZ showed no significant increase in apoptosis (SI Appendix Fig S4).

**Figure 1:**
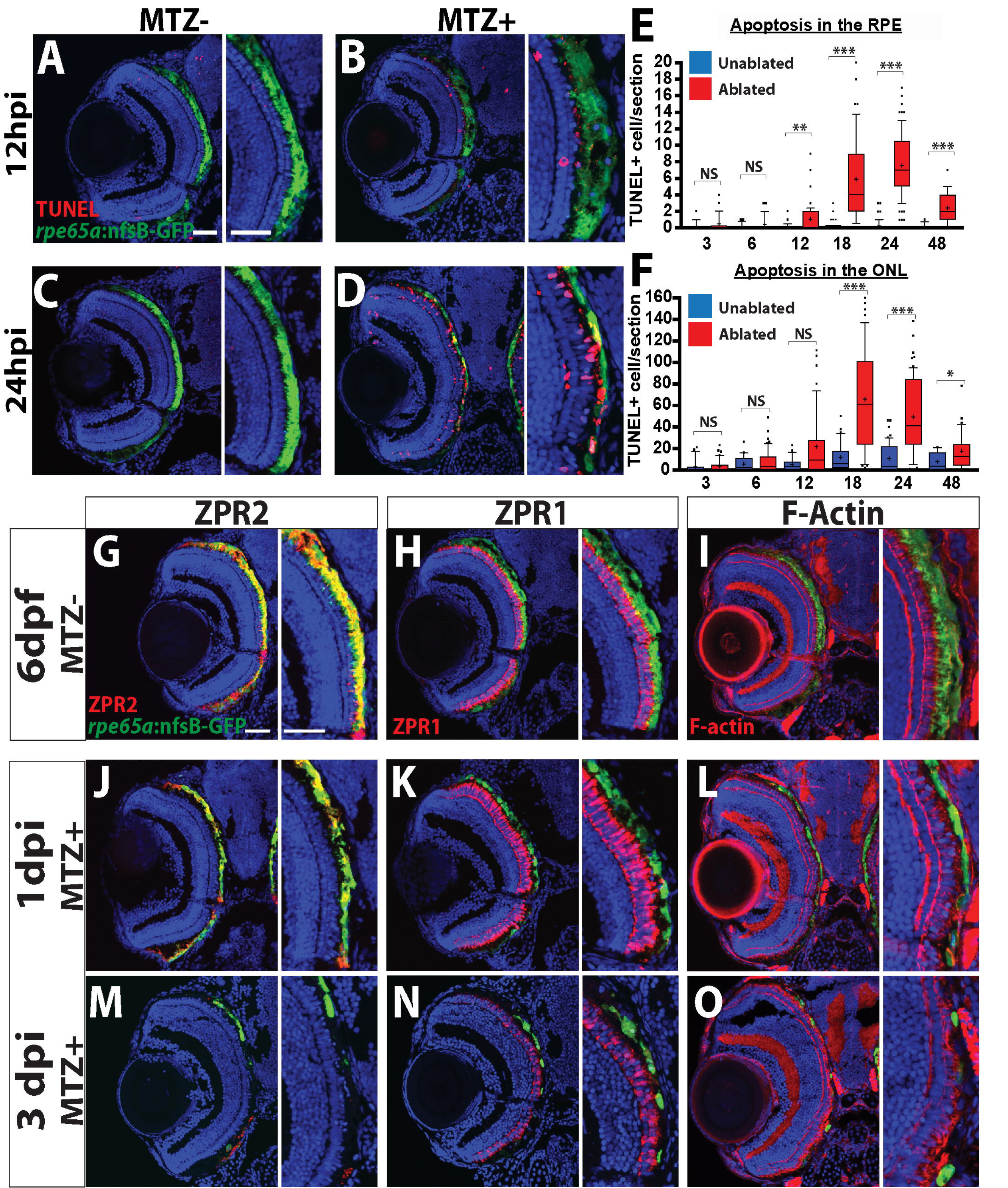
Ablation of the RPE leads to degeneration of underlying photoreceptors. (A-D) Transverse cryosections stained for TUNEL (red). Compared to untreated (A,C) larvae, ablated RPE were disrupted by 12hpi (B), and TUNEL^+^ cells appeared throughout the RPE and ONL at 24hpi (D). (E, F) Quantification of TUNEL^+^ cells/section in the RPE (E) and ONL (F) revealed a significant increase in the RPE by 12hpi and in the ONL by 18hpi. Significance determined using Mann-Whitney U test. * p≤0.05, ** p<0.005, *** p<0.0005. (G-I) Transverse sections of unablated 6dpf larvae stained for ZPR2 (G), ZPR1 (H), and F-Actin (I) (red). By 1dpi, ZPR2 is disrupted in a similar manner to GFP (J), and PR morphology becomes disorganized (K,L). By 3dpi, ZPR2 signal is absent from the injury site (M) and PR morphology is notably degraded (N,O). Green=GFP, blue=nuclei. Dorsal is up and distal is left. Scale bar = 40μm.

To characterize degeneration further, RPE-ablated larvae were stained with markers for RPE (ZPR2) (38), red/green cone arrestin (ZPR1) (39), and F-actin (phalloidin) (Fig 1G-O). In unablated larvae, *rpe65a*:nfsB-GFP was expressed in mature RPE while the ZPR2 signal extended further into the periphery, labeling both mature GFP^+^ RPE and less-mature GFP^−^RPE closer to the ciliary marginal zone (CMZ) (Fig 1G). Between 1 and 3 days post injury (dpi), changes to ZPR2 staining recapitulated disruption of GFP^+^ RPE, including degeneration of the RPE cell body and loss of apical microvilli. ZPR1^+^ cones also began degenerating at 1dpi (Fig 1K), and F-actin bundles in POS became more diffuse and lost their perpendicular orientation (Fig 1L). By 3dpi, both GFP and ZPR2 signals were absent from the central RPE, confirming RPE degeneration in the central injury site (Fig 1M). PR degeneration in the central retina also peaked at this time, displaying aberrant cone morphology (Fig 1N), and significant degeneration of POS throughout the injury site (Fig 1O). Despite rigorous screening, some variability in ablation severity was observed, likely from variations in transgene expression and ablation efficiency. Only larvae with high levels of GFP signal disruption (severe ablation) were utilized in subsequent experiments. In severely ablated larvae, ablation-mediated degeneration reliably peaked between 2-3dpi (i.e. Fig 1).

Immunohistochemical data strongly supported RPE and PR degeneration following ablation and this was confirmed by transmission electron microscopy (TEM) (Figs 2, SI Appendix S5). In unablated larvae, central RPE cells contained pigmented melanosomes (Fig 2A, S5C), formed tight junctions (TJ) with adjacent RPE cells (Fig S5C), extended melanosome-containing apical processes to interdigitate with POS (Fig SI Appendix S5A), and possessed phagosomes containing phagocytosed POS discs (Fig SI Appendix S5C). The PR layer was laminated and contained readily identifiable cone outer segments (cOS) and rod outer segments (rOS) (Fig 2A). Analysis of ablated larvae at 3dpi revealed severe degeneration of the RPE, which was occupied by debris—much of it likely consisting of degrading POS, which were either distributed throughout the injury site or collected in membrane-enclosed structures that may be macrophages (Fig 2B, SI Appendix S5B). Bruch’s membrane (BM) underlying ablated RPE was also significantly thinner than control (Fig 2C-E, p<0.0001) and contained obvious gaps (Fig 2D). Consistent with defects detected by histology (Fig 1), the PR layer of ablated larvae was severely degenerated, showing reduced size and integrity of POS, and containing degenerated POS and other cellular debris (Fig 2B, SI Appendix S3B,D). Lamination of PR nuclei and ribbon synapse (RS) morphology were disrupted in the central injury site, suggesting that PR connectivity to bipolar cells degraded following RPE ablation (SI Appendix Fig S5E,F).

**Figure 2:**
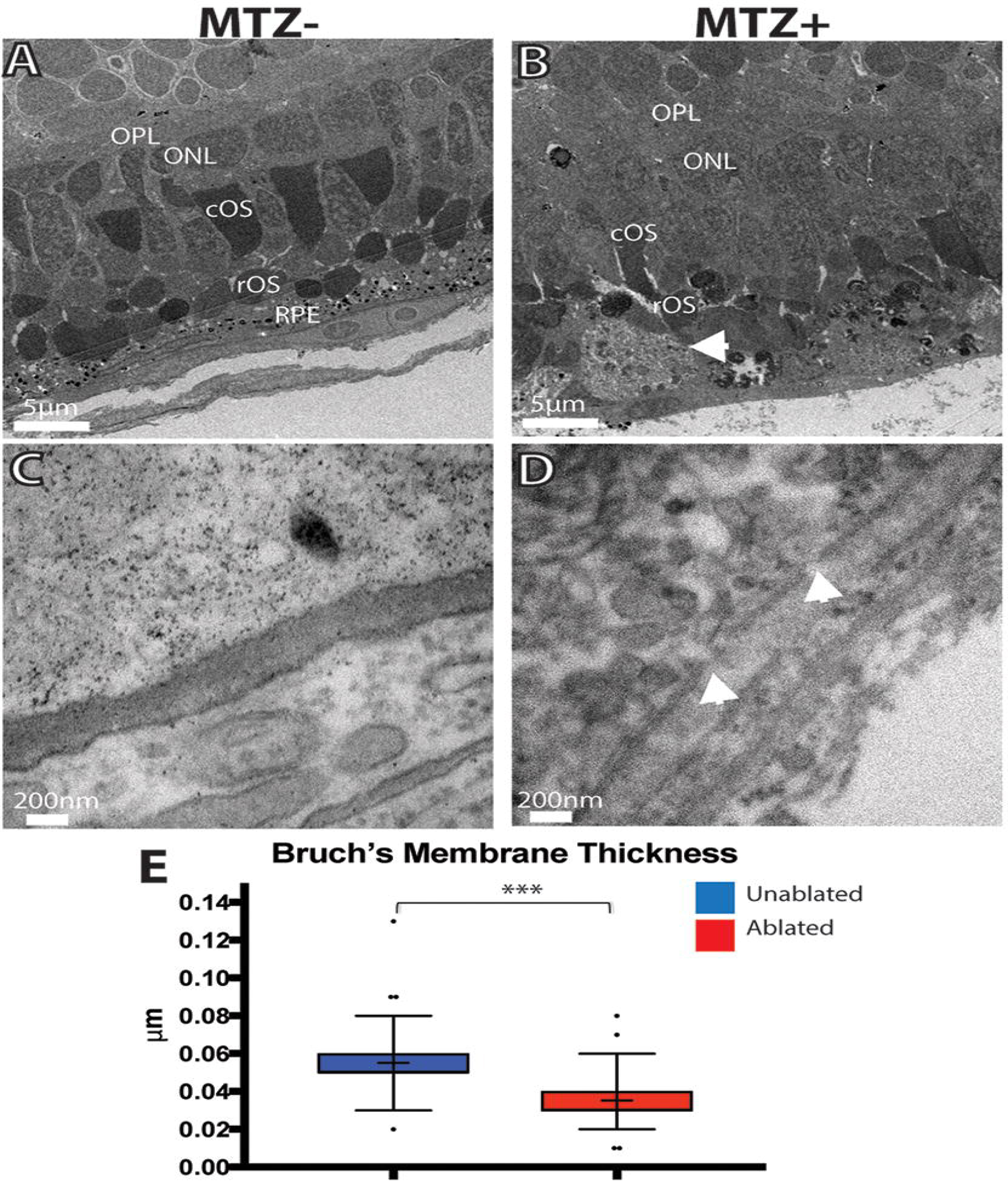
TEM analysis of ablated retina and RPE. (A,C) Unablated 8dpf larvae. (B,D) 3dpi larvae. (A,B) Ablated larvae possess significant degeneration within the ONL and RPE. Aggregates of debris are notable in ablated eyes (arrow, B). Magnification of BM in unablated (C) and ablated (D) larvae shows BM degeneration and obvious gaps perforations (arrows, D). (E) Quantification of BM thickness. Student’s T-test reveals that BM thickness is significantly decreased in ablated larvae *** p<0.0005. (MTZ-n=3 eyes, 81 measurements; MTZ+ n=4 eyes, 108 measurements)

Visual function of larvae was evaluated by analyzing the optokinetic response (OKR) to determine whether RPE ablation results in vision defects (40) (SI Appendix Fig S6). Ablated and control larvae were exposed to a rotating full-field visual stimulus at 1dpi, 2dpi and 3dpi, and visual responses were recorded (SI Appendix Fig. S6A-C). At 1dpi, ablated larvae exhibited a modest reduction in stimulus tracking gain relative to controls, and this reduction in gain became significant at 2dpi (SI Appendix Fig. S6D, p=0.0055) indicating that visual function was disrupted. By 3dpi, ablated larvae demonstrated a recovery of stimulus tracking gain (SI Appendix Fig. S6C,D). Collectively, these data demonstrate that ablation of large swathes of mature RPE in *rpe65a*:nfsB-GFP transgenics results in the rapid degeneration of underlying PRs and BM, and a loss of visual function, defects reminiscent of late-stage atrophic AMD (10).

### Regeneration of the RPE

As discussed above, a subset of RPE cells possess a latent ability to proliferate *in vitro* (30) and various degrees of RPE repair have been documented, but in no system is the RPE able to recover a functional monolayer following a large injury. Zebrafish possess a remarkable ability to regenerate a multitude of tissues, but it is unknown if they can regenerate RPE. Thus, we analyzed the regenerative capacity of ablated larvae at 4, 6, 7, and 14dpi with ZPR2, ZPR1, and phalloidin (Fig 3). At 4dpi, ZPR2^+^ cells extended into the injury site (Fig 3D) and RPE pigmentation significantly increased compared to 2dpi levels (Fig S2B,G), suggesting that RPE cells have begun to regenerate. Although ZPR1^+^ cones and POS remained degenerated in the central ablation site, morphologically normal ZPR1^+^ cones reappeared in the periphery, and these were always in direct apposition to regenerated GFP^+^ RPE (Fig 3E,F, SI Appendix Fig S7). At 6dpi, morphologically normal GFP^+^/ZPR2^+^ RPE cells nearly filled the injury site (Fig 3G), and PR morphology recovery occurred in a similar pattern (Fig 3H,I, SI Appendix Fig S7). Interestingly, ZPR2^+^/GFP^−^ cells always appeared at the advancing tip of the regenerating monolayer (Fig 3G). While the *rpe65a:*nfsB-GFP transgene is expressed specifically in mature RPE cells, ZPR2 labels less-mature RPE, suggesting that these ZPR2^+^/GFP^−^ cells are RPE cells that have not yet fully differentiated. By 7dpi, the injury site was populated by ZPR2^+^ RPE cells (Fig 3J). Although ZPR1-labeled cones remained disorganized in the central retina at 7dpi (Fig 3K), POS architecture began to improve (Fig 3L). By 14dpi, ZPR2^+^/GFP^+^ cells populated the entire RPE layer, and these displayed proper RPE cell morphology (Fig 3M,N). While ZPR1^+^ cones also regenerated, ONL disorganization persisted, particularly in the injury site, where cones failed to align perpendicularly to the RPE (Fig 3N). 7 months post ablation, we observed recovered RPE that were morphologically similar to unablated siblings (SI Appendix Fig S8).

**Figure 3:**
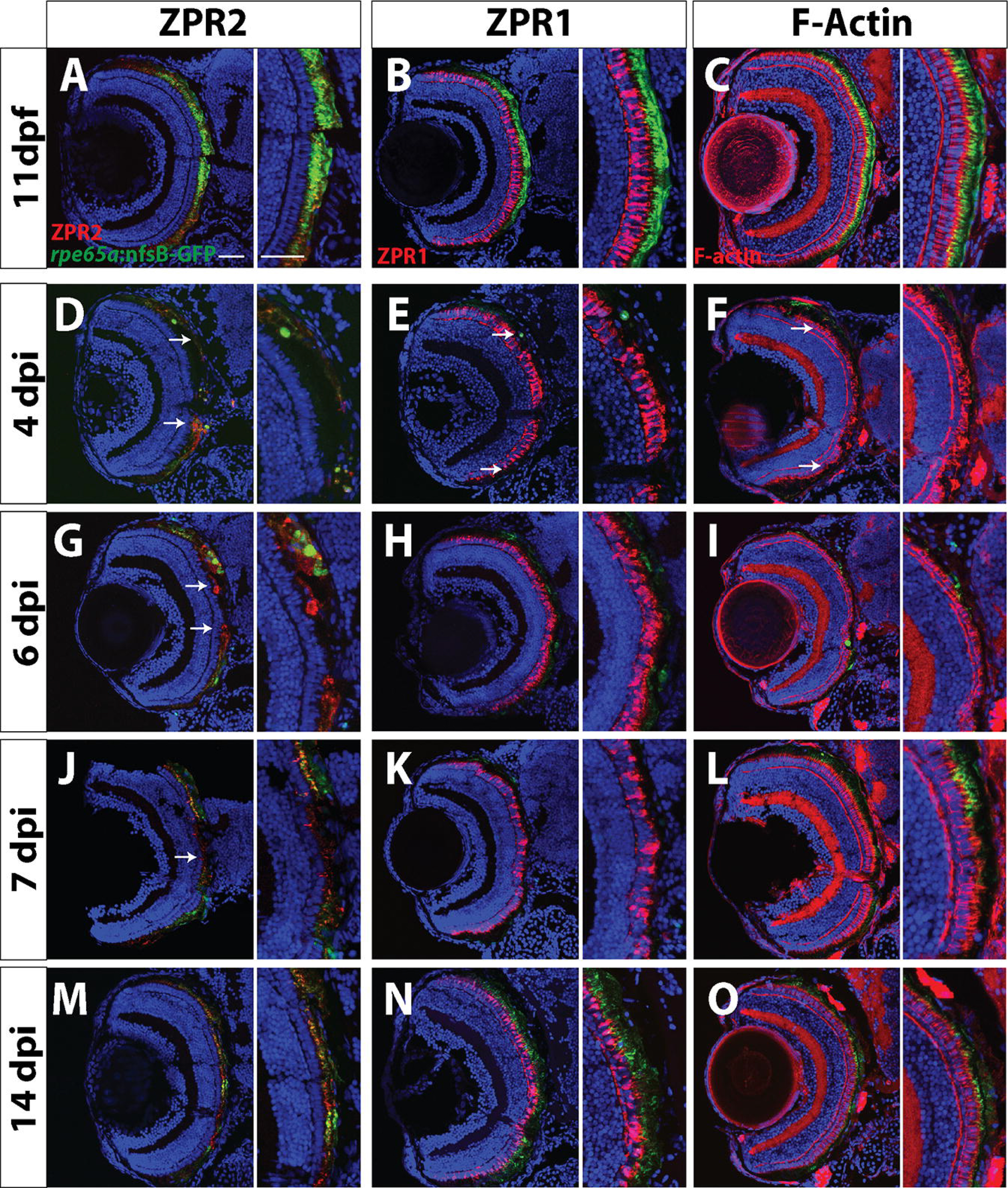
RPE regeneration initiates in the periphery and proceeds inward. Transverse sections of unablated larvae stained for ZPR2 (A), ZPR1 (B) and F-Actin (C) at 11dpf. Ablated eyes stained for ZPR2 (D,G,J,M), ZPR1 (E,H,K,N), and Phalloidin (F,I,L,O) at 4, 6, 7 and 14dpi. Green=GFP, blue=nuclei, red=marker. GFP^+^ RPE appears in the periphery by 4dpi (marked by arrows in D-F). As regeneration proceeds, GFP^+^ RPE extends further toward the eye center, and the leading tip of the regenerated monolayer often consists of both immature and mature RPE (ZPR2^+^/GFP^−^ cells in G). PR morphology appears to recover in the periphery proximal to regenerated RPE. By 7dpi, ZPR2^+^ RPE is present throughout the RPE (J), and PR morphology begins to recover in the central injury site (K,L). By 14dpi, mature GFP^+^/ZPR2^+^ RPE cells are present throughout the RPE (M), and PR morphology further improves in the central retina (N,O). Dorsal is up and distal is left. Scale bar = 40μm.

Regeneration appeared to proceed in a periphery-to-center fashion in fixed samples. To test this *in vivo*, we utilized optical coherence tomography (OCT) to quantify the spatial and temporal dynamics of RPE degeneration and regeneration in individual larva over time. When intact, the RPE in OCT images presents as a bright line due to the density and pigment present; in ablated eyes, the intensity of the signal decreases as a result of tissue disruption (Fig. 4A,B). RPE signal intensity (backscatter) can be quantified by determining the pixel intensity at each position of the RPE; here, we quantified the intensity from the optic nerve to the dorsal periphery, and examined changes in intensity in individual larvae over time. This analysis revealed that backscatter was significantly decreased in ablated larvae compared to controls in the central-most three quintiles of the RPE at 1dpi, and that all but the central-most quintile recovered to unablated levels by 5dpi (Fig 4C, p<0.0001). These results demonstrate that RPE recovery occurs in a peripheral-to-central manner.

**Figure 4:**
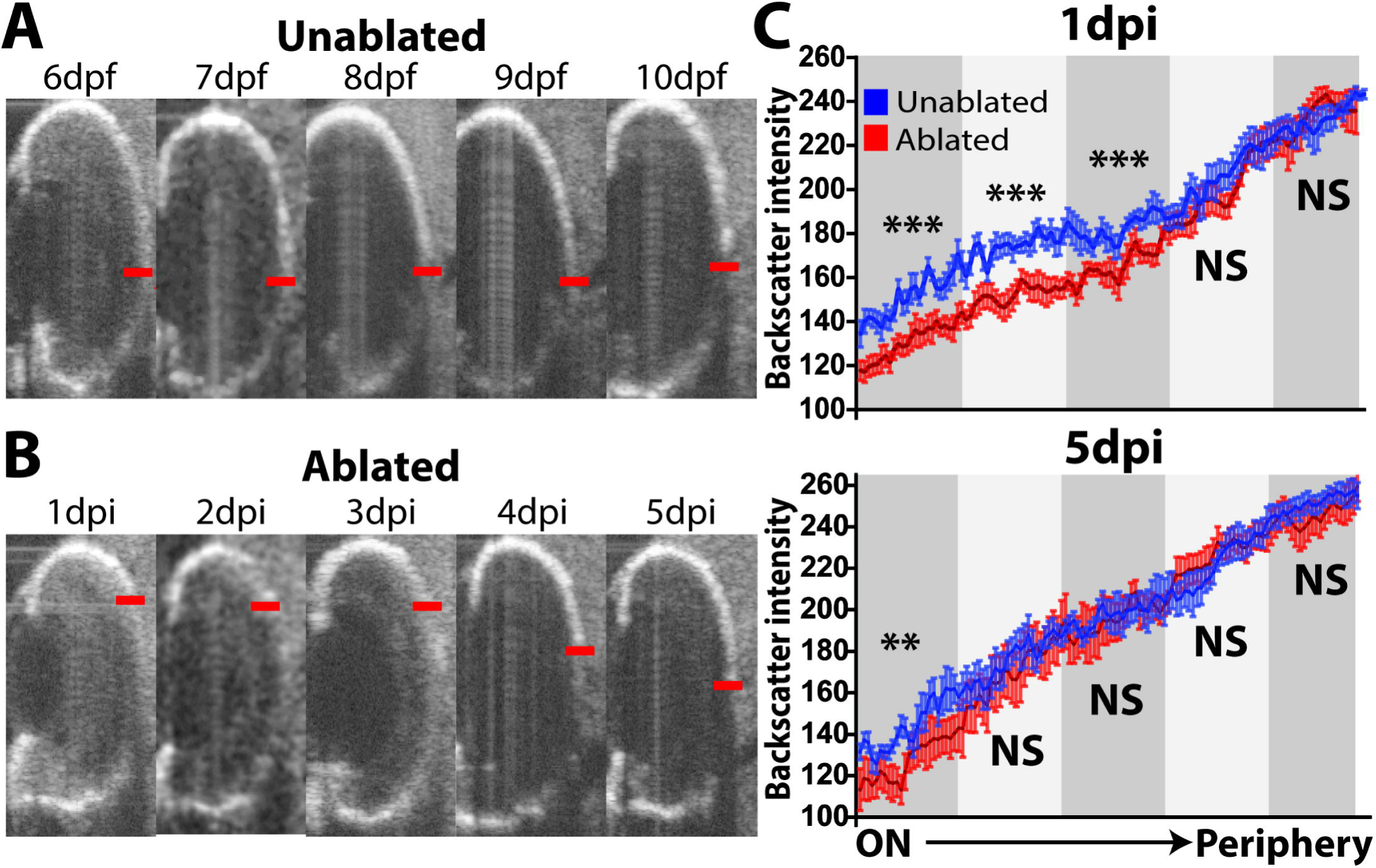
Longitudinal OCT analysis of RPE regeneration. (A-C) Representative time series from a single (A) unablated and (B) ablated larva. Central edge of maximal RPE intensity marked with red line. (C) Quantification of the RPE signal (backscatter) from the dorsal periphery to optic nerve across unablated (blue) and ablated (red) larvae at 1dpi and 5dpi (error bars=SEM). The measured RPE was divided into quintiles, and the area under the curve within each quintile was measured. At 1dpi, backscatter intensity in ablated RPE is significantly below unablated intensity in the 3 quintiles closest to the optic nerve, while at 5dpi, only the central-most quintile is significantly reduced (MTZ^−^: n=10, MTZ^+^ n=9, Student’s unpaired t test, **p<0.005, ***p<0.0005).

To confirm that regenerated RPE is morphologically normal and functional, TEM was performed on 14dpi larvae, a time at which GFP^+^/ZPR2^+^ RPE cells populate the injury site (Figs 5, SI Appendix S9). Both unablated and ablated RPE contained melanosomes, extended apical processes to interdigitate with POS, and contained phagosomes (Fig 5A,B, SI Appendix S9A-D). In ablated larvae, BM thickness was restored (Fig 5C-E, p=0.3402). However, differences existed between regenerated and unablated RPE: individual RPE cells in the regenerated region appeared to contain more melanosomes and had thicker cell bodies (Fig 5A,B, Fig SI Appendix S9A,B). Consistent with immunohistochemical results, ONL lamination was improved but not completely recovered. Nonetheless, the presence of morphologically normal ribbon synapses in the central injury site suggests recovery of PR connectivity to bipolar cells (Fig SI Appendix S9E,F). Taken together, these data demonstrate that zebrafish are capable of regenerating a functional RPE following widespread ablation, and that regeneration is rapid, occurring within 1-2 weeks post ablation.

**Figure 5:**
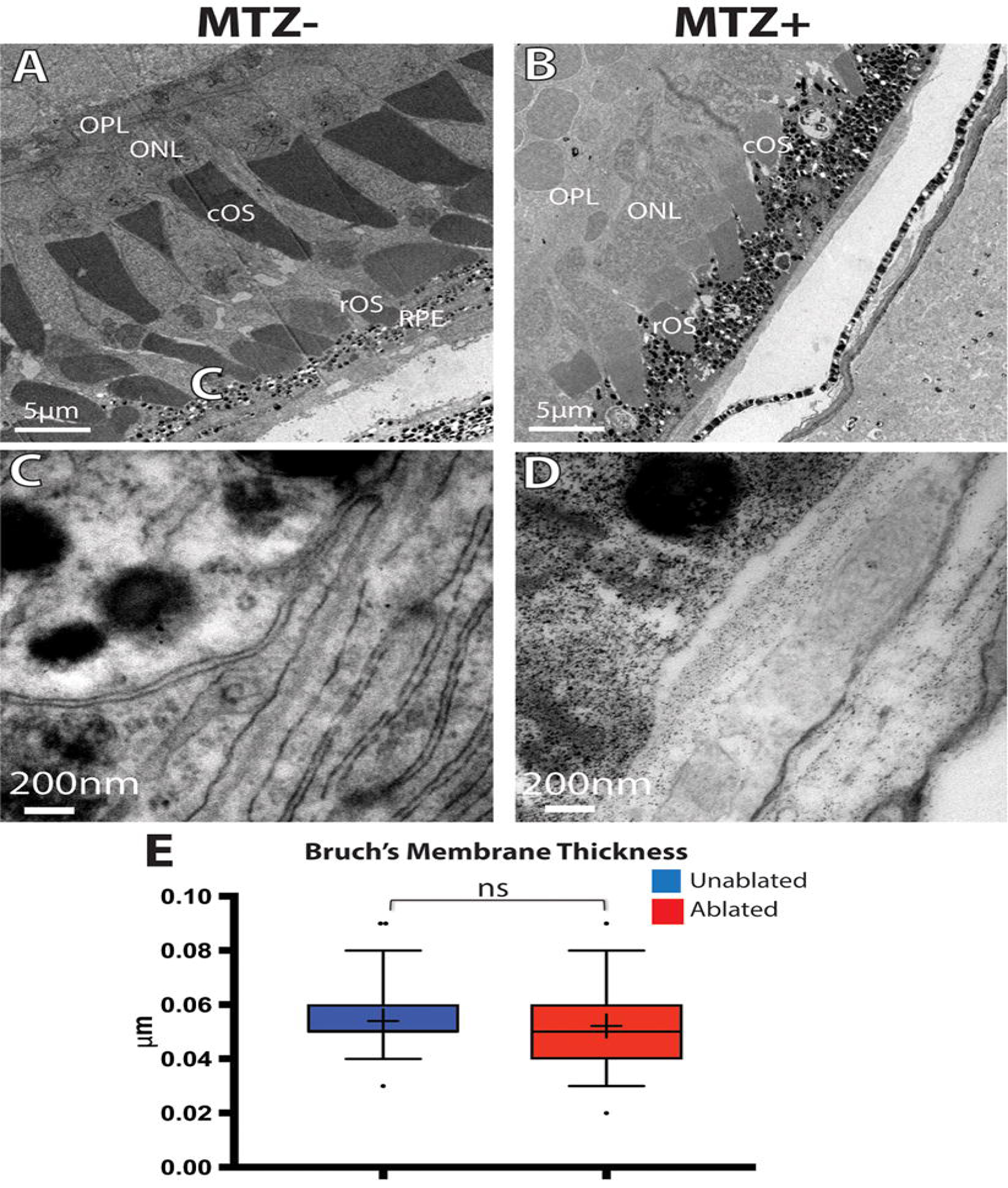
TEM analysis of regenerated RPE. A,C) Unablated 19dpf larvae. (B,C) 14dpi larvae. (A,B) Images of unablated RPE and ONL ultrastructure. Regenerated POS are visible in the ablated ONL, and a regenerated RPE is present (B). Magnification of BM in unablated (C) and ablated (D) larvae reveals the presence of a regenerated BM appears in ablated larvae. (E) Quantification of BM thickness. Student’s T-test reveals that BM thickness is not significantly decreased in ablated larvae * p<0.05. (MTZ^−^ n=3 eyes, 81 measurements; MTZ^+^ n=3, 81 measurements)

In larvae, the eye undergoes significant growth, making it possible that RPE regeneration is the result of a permissive growth environment rather than an ability of the RPE to regenerate *per se*. Thus, we assessed whether RPE regeneration also occurs in the adult eye. Transgene expression in adults is restricted to mature central RPE as it is in larvae (Fig S10A). At 2dpi, there were clear signs of RPE degeneration that mirrored those in RPE-ablated larvae, including disruption of cell body cohesion and degeneration of apical processes, as indicated by the aberrant colocalization of ZPR2 and GFP in the photoreceptor layer (SI Appendix Fig S10C,D). By 14dpi, adults showed signs of RPE regeneration in the peripheral injury site, such as recovery of contiguous GFP^+^/ZPR2^+^ RPE (SI Appendix Fig S10E, arrows) and apical polarization of ZPR2-localization (SI Appendix Fig S10F). By 28dpi, GFP^+^/ZPR2^+^ RPE extended to the injury site (SI Appendix Fig S10G, arrows) and ZPR2 staining showed restoration of RPE apical processes (SI Appendix Fig S10G,H). Taken together, these results demonstrate that the adult zebrafish is also capable of regenerating the RPE, and in a similar peripheral-to-central manner as in larvae. Given these similarities and the technical advantages of using larvae over adults (e.g. comparatively rapid regeneration, access to a large number of samples, feasibility of *in vivo* imaging, and utility in high-throughput drug screens), we focused further efforts towards on characterizing the mechanisms underlying RPE regeneration in larvae.

### Cellular Dynamics Underlying RPE Regeneration

The rate and peripheral-to-central pattern of RPE regeneration suggest that regeneration is driven by cell proliferation, and not simply an expansion of RPE, a response noted in several systems after small injuries (19, 26, 41). Proliferative cells are a major component of regeneration in diverse tissues, and they often derive from a resident pool of progenitor cells (42, 43), or from differentiated cells that are stimulated to respond to injury (33, 44). Moreover, RPE proliferation results from loss of BM contact in several injury contexts, and pathologically, during PVR (19, 25, 45). That we observed BM degeneration throughout the large injury site makes it highly unlikely that peripheral cells could expand to cover the entire span of lost RPE. Thus, we hypothesized that uninjured peripheral RPE cells respond to injury by dedifferentiating and proliferating to replace lost tissue.

To test this hypothesis, we exposed larvae to 2-hour pulses of EdU, dissected eyes, and stained for ZPR2/EdU, imaging them to acquire *en face* views of the RPE (Fig 6). 24-hour BrdU incorporation assays were also performed to characterize the total numbers of proliferative cells within the RPE throughout regeneration (Fig S11). Proliferative cells appeared in the RPE as early as 1dpi, largely appearing immediately adjacent to the CMZ or in the center of the ablation site (Figs 6F,N; SI Appendix S11G,U, p<0.0001). Between 2-3dpi, more proliferating cells localized to the center of the eye, within the injury site (Figs 6G,H; SI Appendix S11I), and the number of proliferative cells in the RPE peaked between 3-4dpi (SI Appendix Fig S11J,U). During this period, proliferative cells populated much of the central eye in ablated larvae, with many localizing adjacent to or within the injury site (Figs 6H,I; SI Appendix S11J,Q-T). In contrast, unablated eyes showed GFP^+^ RPE cells throughout the central RPE (Fig 6C,D; SI Appendix S11D,M-P) and sparse BrdU^+^ nuclei (SI Appendix Fig S11M-P). As regeneration continued, GFP^+^ RPE cells appeared closer to the center of the injury site and the number of proliferative cells in the RPE layer decreased (Figs 6I,J; SI Appendix S11L,U), with most remaining proliferative cells localizing to the injury site. As expected, BrdU^+^ cells were also observed in the retina, and these are likely Müller glia-derived progenitor cells (MGPCs) generated in response to PR degeneration (33). Quantification of BrdU^+^ cells in the central retina demonstrated that the kinetics of retinal regeneration largely overlapped that of the RPE (SI Appendix Fig S11V).

**Figure 6:**
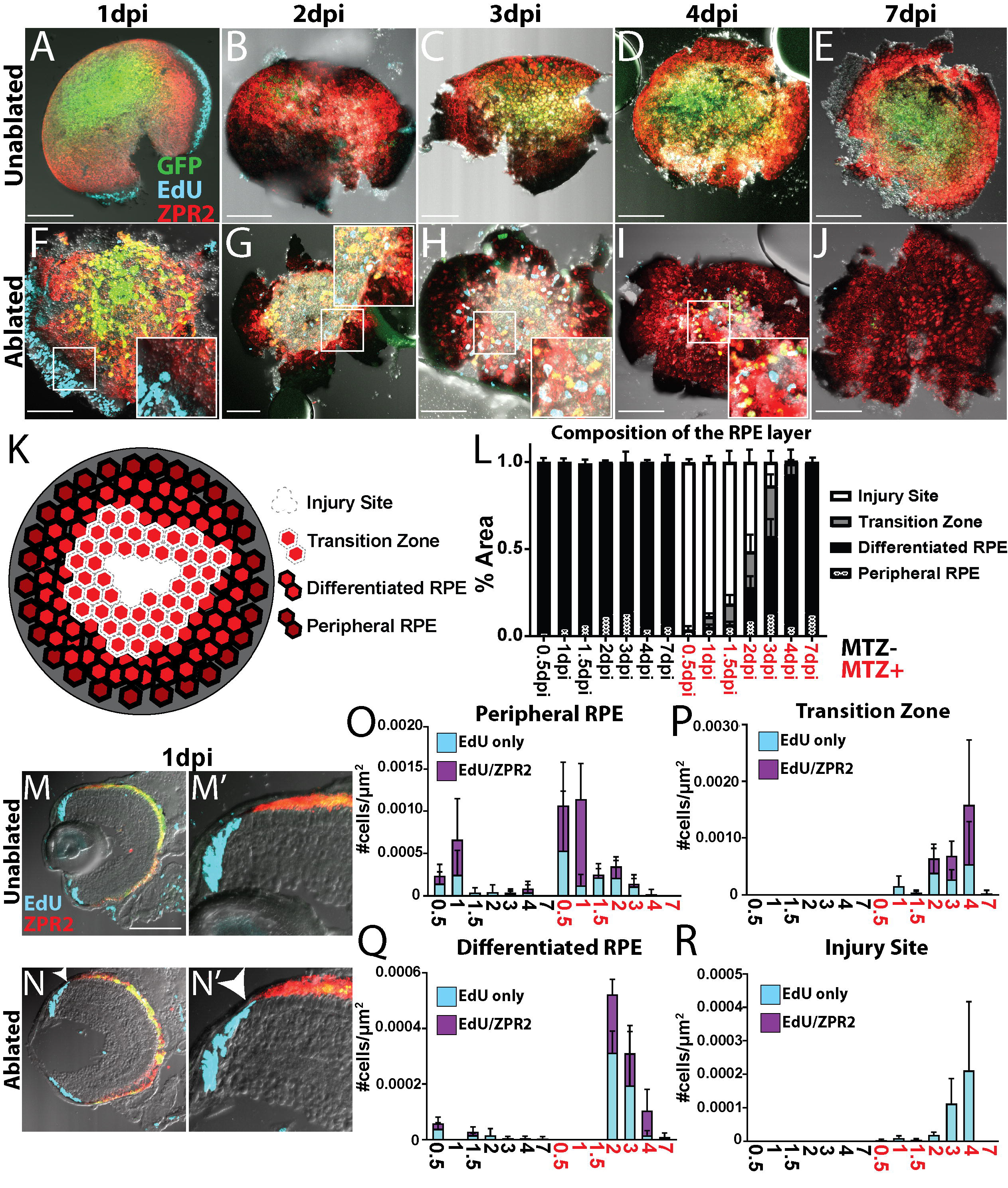
Wholemount analysis of RPE cell proliferation and regeneration. (A-J) *en face* wholemount images of unablated (A-E) and ablated (F-J) larvae exposed to 2-hour EdU pulses immediately before fixation and staining for ZPR2 at various time points post injury. (K) Division of RPE into 4 domains, cartoon. (L) Quantification of the area of RPE comprised by each domain during regeneration. (M-N) Transverse cryosections of unablated (M) and ablated (N) eyes at 1dpi. Magnified insets (M’,N’) reveal the presence of EdU^+^ cells in the RPE periphery in ablated retinae (N’ arrowhead). (O-R) Quantification of the density of EdU cells throughout regeneration in the peripheral RPE (O), Transition Zone (P), differentiated RPE (Q) and Injury Site (R).

To quantify the spatial dynamics of RPE regeneration and RPE cell proliferation, the RPE of each eye was divided into four regions based on pigmentation level, ZPR2 expression, and location: (1) peripheral RPE (pRPE; pigmented, dimly ZPR2^+^), (2) differentiated RPE (dRPE; pigmented, ZPR2^+^), (3) transition zone (TZ; lightly pigmented, ZPR2^+^) consisting of incompletely differentiated RPE extending into the injury site, and (4) injury site (IS; unpigmented, ZPR2^−^) (Fig 6K). Using these criteria to quantify RPE layer composition over time, this analysis confirmed that a large proportion of the RPE degenerates rapidly after ablation, and it revealed that the TZ appears as soon as 1dpi, with dRPE reappearing in the periphery at 2dpi. Regeneration of a pigmented ZPR2^+^ RPE was completed by 7dpi (Fig 6L). During regeneration, ZPR2^+^ TZ cells always extended further centrally than dRPE, and the proportion of the TZ at each time point correlated with the proportion of the dRPE added to the RPE the following time point, strongly suggesting that TZ cells become dRPE. Quantification of the density of EdU^+^ and EdU^+^/ZPR2^+^ cells within each region revealed that there are more EdU^+^/ZPR2^+^ cells in the pRPE in ablated larvae at 0.5dpi and 1dpi, and though this increase did not achieve significance (Fig 6O, p=0.076 and p=0.078, respectively), cryosections of larvae at 1dpi showed peripheral EdU^+^ZPR2^+^ cells similar to those observed after BrdU exposure (compare Fig 6M,N to Fig S11G, arrow). During these early time points, EdU^+^/ZPR2^+^ cells were largely restricted to the peripheral retina, with only a few EdU^+^ cells appearing in the IS (Fig 6R). During intermediate time points, when dRPE reappears and the TZ extends centrally, EdU^+^/ZPR2^+^ cells were present in both dRPE and the TZ. As regeneration proceeded, the density of EdU^+^/ZPR2^+^ in dRPE decreased, while increasing in the TZ (Fig 6P,Q). By 4dpi, proliferative cells were largely restricted to the center of the eye (Fig 6I), and the majority of the remaining proliferative cells were located either in the IS or TZ. Interestingly, the TZ and dRPE contained an even mix of EdU^+^ and EdU^+^/ZPR2^+^ cells, which may suggest that some differentiated RPE cells remain proliferative in this region and continue to generate new EdU^+^/ZPR2^−^ cells that later enter the TZ and differentiate. Taken together, these results support a model in which peripheral RPE responds to injury by proliferating, that proliferative RPE cells move into the IS, and that proliferation continues within newly-generated RPE adjacent to the injury site until the lesion is repaired.

To determine whether early-proliferative cells enter the injury site and continue proliferating, we pulsed ablated larvae with BrdU between 0-1dpi, and with EdU at 3dpi before fixation and analysis (Fig 7A-B). Transverse sections revealed a significant increase of BrdU^+^/EdU^+^ cells and BrdU^+^ cells within the RPE layer in ablated fish (Fig 7C, p<0.0001). Interestingly, BrdU^+^/EdU^+^ cells often appeared at the interface between pigmented RPE and the unpigmented injury site, and some appeared to be pigmented (Fig 7B”). We next sought to determine whether early-proliferative cells ultimately integrate into the regenerated RPE. To do this, we exposed ablated larvae to BrdU between 0-1dpi and fixed them at 7dpi for analysis (Fig 7D,E). Transverse sections revealed a significant increase of BrdU^+^ cells within the RPE (Fig 7E,J, p<0.0001). These data suggest that early-proliferative cells enter the injury site at the leading edge of the regenerating RPE and either continue proliferating or give rise to proliferative cells there. Were this the case, we hypothesized that early-proliferative cells would integrate into both the peripheral and central RPE layer, while later-proliferative cells would only form RPE within the central RPE. To test this, we pulsed ablated larvae with BrdU at 3-4dpi or 5-6dpi before fixing at 7dpi (Fig 7F-I). BrdU^+^ cells were distributed throughout the RPE after a 0-1dpi pulse, but became more restricted to the central RPE after 3-4dpi and 5-6dpi pulses (Fig 7K). Finally, to determine whether early-labeled proliferative cells ultimately differentiate into RPE by 7dpi, we quantified the number and centrality of BrdU^+^/GFP^+^ cells in 7dpi larvae that had been pulsed with BrdU between 0-1dpi (SI Appendix Fig S12). Our analysis revealed that significantly more BrdU^+^ cells in the RPE were GFP^+^ than GFP^−^ (SI Appendix Fig S12B, p=0.0005), and that BrdU^+^/GFP^+^ cells preferentially integrated toward the center (SI Appendix Fig S12C, p<0.0001).

**Figure 7:**
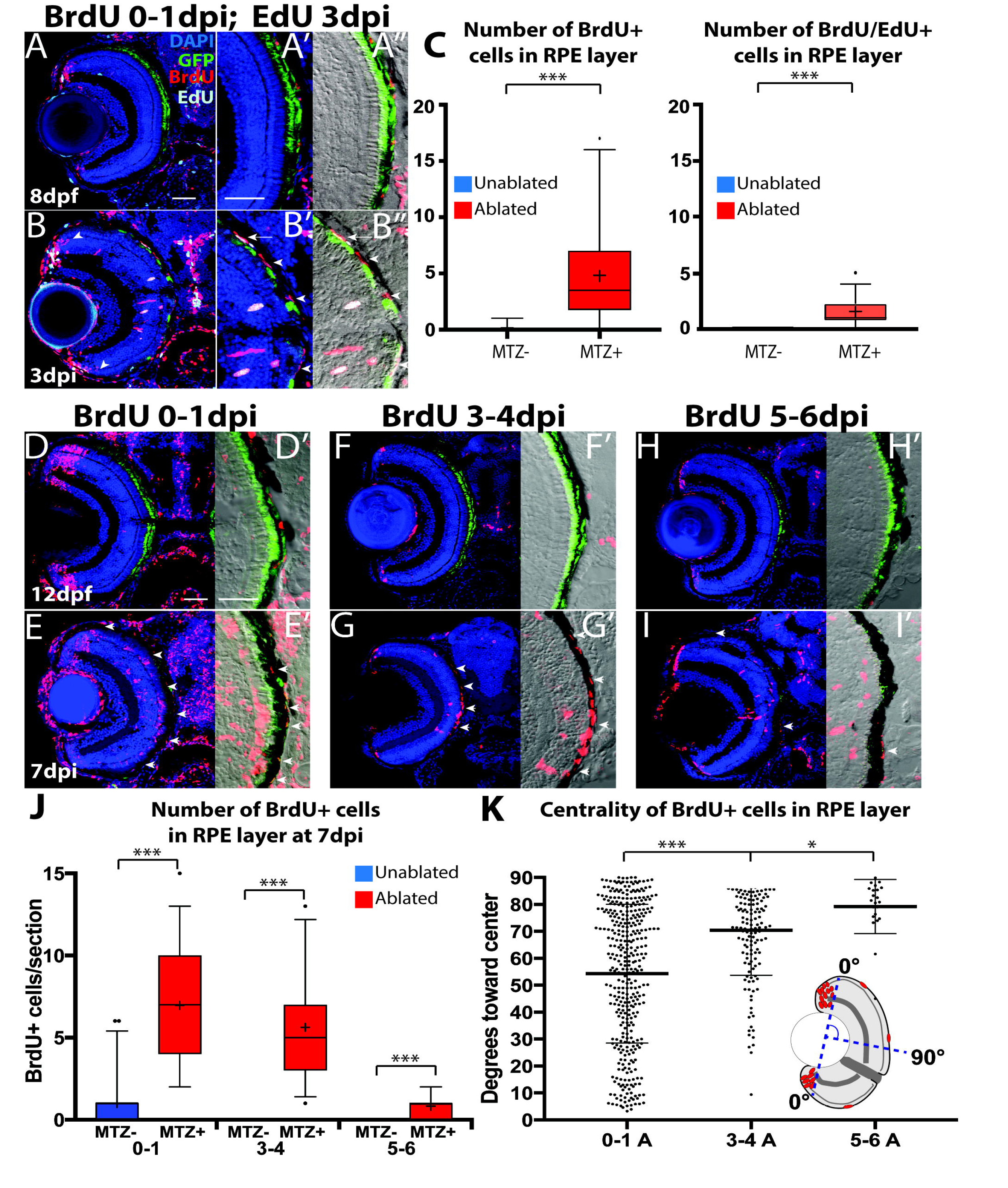
Proliferative RPE contributes to the regenerated RPE monolayer. (A-A”) Transverse sections from unablated larvae exposed to BrdU from 5-6dpf and pulsed with EdU for 2 hours before fixation at 8dpf. (B-B”) Transverse sections of ablated larvae exposed to BrdU from 0-1dpi and pulsed with EdU for 2 hours before fixation at 3dpi. (A’,B’) Magnified inset of BrdU/EdU. (A”,B”) Magnified inset of BrdU/EdU and DIC. Arrowheads in (B) highlight BrdU^+^ PRs that have integrated into the ONL. Arrow in (B’,B”) highlights a proliferative RPE cell, and arrowheads highlight unpigmented, previously-proliferative RPE-like cell in the injury site. (C) Quantification of BrdU/EdU^+^ and BrdU^+^ nuclei in the injury site. (D,E) Larvae exposed to BrdU 0-1dpi and fixed at 7dpi. (F-G) Larvae exposed to BrdU 3-4dpi and fixed at 7dpi. (H,I) Larvae exposed to BrdU 5-6dpi and fixed at 7dpi. (J) Quantification of BrdU^+^ cells per section. (K) Quantification of the location of individual BrdU^+^ cells relative to the center of the RPE. The line indicates the average location of BrdU^+^ cells, and the whiskers indicate standard deviation. Mann-Whitney U Test, * p<0.05, ** p<0.005, *** p<0.0005. Dorsal is up and distal is left. Scale bar = 40um.

### Wnt pathway activity is required for RPE regeneration

Previous studies have identified Wnt signaling as a regulator of regeneration in multiple contexts (46–49), including the retina (50–52). Thus, we examined Wnt signaling to begin to gain mechanistic insight into the molecular mechanisms underlying RPE regeneration. To assess Wnt pathway activity after RPE ablation, we examined expression of the Wnt target gene, *lef1* (53). *lef1* was upregulated in ablated larvae at 1dpi (Fig 8B), but not in unablated siblings (Fig. 8A). Closer analysis of *lef1* expression revealed transcripts distributed in and adjacent to the RPE (Fig 8B’), suggesting the Wnt pathway is activated post-ablation. We next utilized the Wnt pathway inhibitor IWR-1 (54) to determine if Wnt activity is required for RPE regeneration. Larvae were pre-treated 24 hours prior to ablation (4dpf/-1dpi) with 15μM IWR-1 or with a vehicle control (0.06% DMSO) and kept in drug or vehicle until fixation at 4dpi (the time at which peak proliferation is observed in the RPE (SI Appendix Fig S11U). Quantification of BrdU^+^ cells/section revealed a significant decrease in proliferation in IWR-1-treated RPE when compared to controls (Fig 8E-G, p<0.0005). Further, there was a noticeable lapse in recovery of a pigmented monolayer in IWR-1-treated larvae (Fig. 8I, *arrowheads*) relative to DMSO controls (Fig 8H). ZPR2 staining overlapped with pigmented RPE in both ablated DMSO-(SI Appendix Fig S13A) and IWR-1-treated (SI Appendix Fig S13B) larvae, indicating the lapse in pigment recovery was not a pigmentation deficiency, but a failure of RPE to regenerate. Quantification of percent RPE recovery indeed showed a significant decrease in the IWR-1-treated larvae (Fig 8J, p<0.0005).

**Figure 8:**
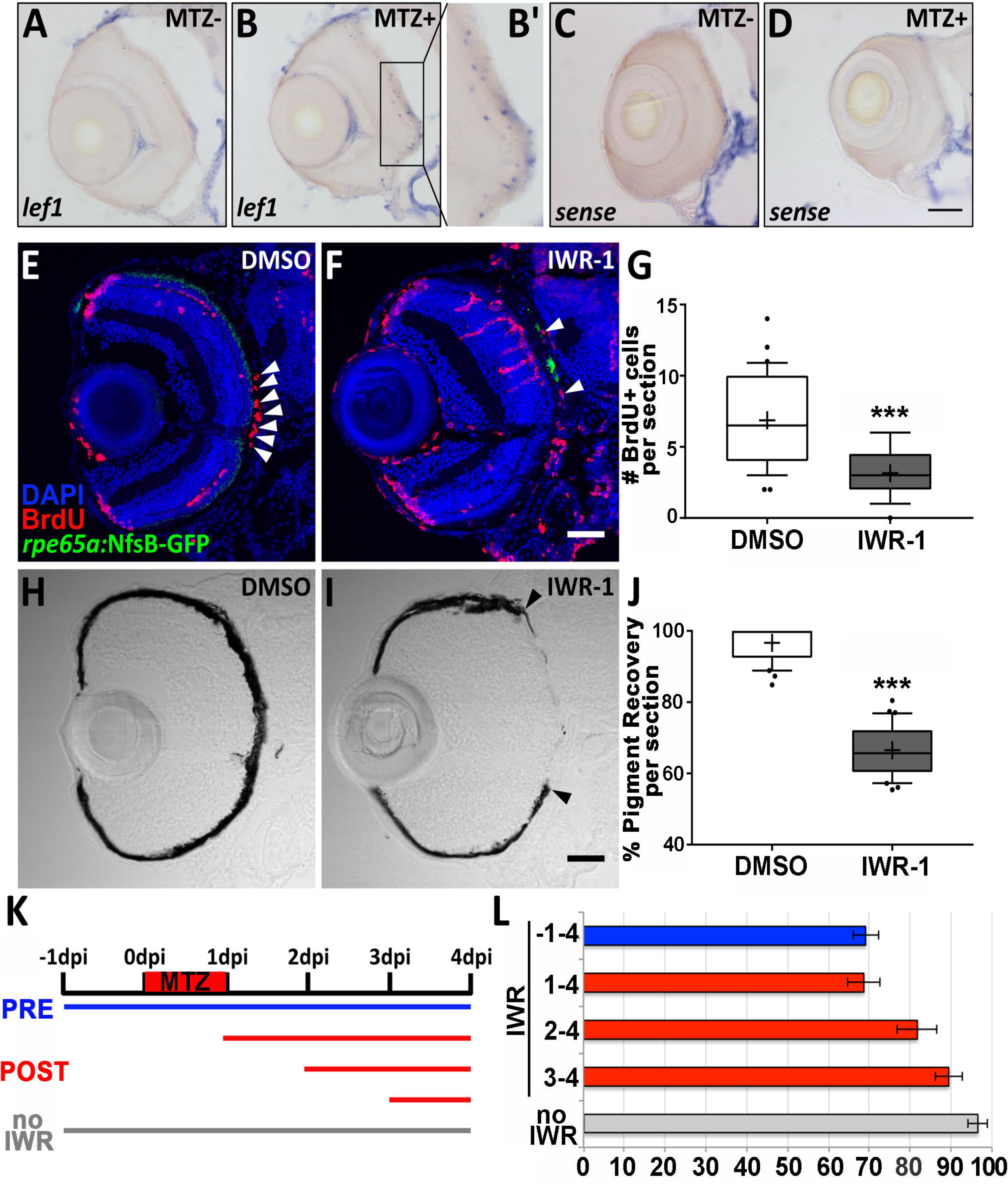
Wnt pathway inhibition impairs RPE regeneration. (A-D) Transverse sections of *lef1* or sense RNA expression in unablated 6dpf (MTZ^−^) and ablated 1dpi (MTZ^+^) larvae. *lef1* is detected in and around the RPE in MTZ^+^ (B’) but not MTZ^−^ larvae. *lef1: n*>*5*; *lef1 sense*: *n*=*4*. (E-J) Transverse sections of 4dpi ablated DMSO-(E,H; *n*=*10*) and 15μM IWR-1-treated (F,I; *n*=*11*) larvae exposed to a 24-hour pulse of BrdU from 3-4dpi. (E,F) Green=GFP, blue=DNA, red=BrdU; *white arrowheads* highlight BrdU^+^ cells in the RPE. (G) Quantification of BrdU^+^ cells/section reveals that IWR-1 treatment significantly decreases the number of proliferative cells in the RPE at 4dpi (Student’s unpaired t-test, *** p<0.0005). Brightfield images (H,I) and quantification of percent RPE recovery/section (J) shows a significant delay in recovery of a pigmented monolayer in IWR-1 treated larvae (Student’s unpaired t-test, *** p<0.0005). (I) *Black arrowheads* indicate the central-most edge of the regenerating RPE. (K) Schematic representation of the 15μM IWR-1 window experiment showing the time course of pre-injury IWR-1 treatment *(blue)*, post-injury IWR-1 treatment *(red)*, and no IWR-1 treatment *(gray)* groups. (L) Quantification of percent RPE recovery/section indicates that Wnt activation is critical between 1-4dpi, but not between −1-1dpi. Error bars represent standard error of the mean (SEM). Pre-injury IWR-1, *n*=*5*; post-injury IWR-1, *n*=*6* for all treatment groups; no IWR-1, *n*=*6* [See Table S1 for p-values]. Dorsal is up and distal is left. Scale bars = 40μm.

To narrow down a possible time during which Wnt signaling is critical for RPE regeneration, we treated larvae with 15μM IWR-1 pre-(−1-4dpi) or post-(1-4, 2-4, 3-4dpi; Fig. 8K) RPE injury. Although larvae treated longer with IWR-1 tended to display the largest deficits in RPE regeneration, (Fig. 8L), no significant difference was observed in RPE recovery between larvae exposed to IWR-1 between −1-4dpi and 1-4dpi, suggesting that Wnt signaling is not required for the immediate injury response (e.g. from −1-1dpi) (Fig 8L, p=0.88). However, shorter exposures to IWR-1, even during later time points (e.g. 3-4dpi), all elicited significant recovery deficits (Fig 8L, p=0.003), indicating that Wnt activity is likely required for later aspects of RPE regeneration.

## DISCUSSION

The stimulation of endogenous RPE regeneration is an appealing possibility for treating degenerative RPE diseases. However, the development of such a therapy is constrained by the paucity of data regarding the cellular and molecular underpinnings of regeneration. While the mammalian RPE possesses a latent proliferative ability, the process by which RPE cells respond to damage by proliferating and regenerating a functional monolayer, rather than overproliferating, as in PVR, remains largely unknown. The development of an animal model of RPE regeneration following widespread damage is a critical first step towards elucidating the regenerative process. Here, we developed a zebrafish model to ablate mature RPE and assess its regenerative capacity. Remarkably, zebrafish regenerate the RPE within 7-14dpi in larvae, and within 1 month in adults. Regenerated RPE cells appear at the periphery of the injury site soon after ablation, and regeneration proceeds in a periphery-to-center fashion. RPE regeneration involves a robust proliferative response wherein proliferative cells first appear at the periphery of the injury site, later populating it and contributing to the regenerated RPE.

In this model, RPE ablation results in retinal degeneration that resembles late-stage atrophic AMD. While the specific causes of AMD remain unclear, data strongly support that RPE dysfunction and degeneration precede PR loss, and that chronic RPE inflammation contributes to RPE and PR death (55). Mirroring retinal degeneration in atrophic AMD, RPE ablation in zebrafish led to RPE apoptosis, BM and PR degeneration and loss of visual function. In comparison, most RPE injury/regeneration models create small lesions using non cell-specific injury techniques (e.g. debridement or laser photocoagulation (27, 56, 57)), or ablate a diffuse subpopulation of RPE cells via sodium iodate (58–60). In mouse, a genetic RPE ablation system expressing diphtheria toxin in a subpopulation of RPE cells did not cause BM degradation or RPE proliferation (41). Indeed, many RPE injury models preserve an intact BM or spare large regions of RPE. In contrast, our zebrafish model creates a severe and widespread RPE injury, which resembles defects observed in late-stage AMD and thus represents a clinically-relevant starting point for studying RPE regeneration.

Our data provide strong evidence that zebrafish larvae rapidly regenerate the RPE: regenerated RPE cells appear at the periphery of the injury site at 2dpi, and the entire lesion is repaired within 1-2 weeks. More surprising was the recovery of visual function by OKR at 3dpi, well before complete RPE regeneration. The threshold number of PRs required for a positive OKR in zebrafish is unknown. However, since the OKR is elicited by a large-field stimulus, activating PRs throughout the eye, this rapid recovery is likely driven by new CMZ-derived PRs that integrate into the peripheral retina by 3dpi rather than regeneration. Consistent with this hypothesis, we observed CMZ-derived BrdU^+^ PRs in the retinal periphery at 3dpi (Fig 7B). More work is required to determine whether these PRs are responsible for this recovery, or whether PRs adjacent to newly-regenerated RPE in the peripheral injury site also contribute to the OKR.

Mammals largely fail to regenerate a functional RPE monolayer following injury (20, 21). One exception to this is in “super healer” MRL/MpJ mice, which regenerate RPE within ∼30 days after administration of mild doses of sodium iodate that cause central RPE degeneration (61). Beyond this, the mammalian RPE is incapable of regenerating after severe injuries (e.g. (22–24, 26)). RPE in newt (62, 63), *Xenopus* (64), and embryonic chick (65, 66), are capable of transdifferentiating and regenerating the neural retina after traumatic injury. However, studies in these models have focused on the RPE-to-retina transdifferentiation process, and RPE-specific regeneration remains unclear.

Our data support the following model of RPE regeneration (Fig. 9): injury-adjacent RPE cells expand into the injury site, where they encounter degraded BM and proliferate to form daughters that enter the injury site and differentiate into RPE. RPE cells commonly expand to fill territory vacated by lost RPE (19, 41), and contact with a degenerated BM induces RPE proliferation in many contexts (19, 25, 29, 67–69). Supporting this, we found that early-dividing cells (0-1dpi) often appear in the RPE periphery, localize to the injury site during peak phases of regeneration, and ultimately form RPE cells that integrate into the regenerated RPE monolayer. Wholemount analyses indicated that proliferative cells appear in the peripheral RPE soon after injury, and proliferative cells differentiate into RPE in distinct zones: (1) newly differentiated injury-adjacent RPE (dRPE), (2) a Transition zone, containing actively differentiating RPE cells, and (3) the Injury site, which contains proliferative cells that do not yet express RPE markers. Further experiments are necessary to determine whether all injury-adjacent RPE cells are capable of proliferating in response to injury, or if proliferation occurs within a subpopulation. Several lines of evidence suggest the latter possibility, and highlight the important role played by peripheral RPE: in mouse, a subpopulation of mature RPE cells in the periphery remain in the cell cycle and respond to microscopic photocoagulation injuries by proliferating at a higher rate than central RPE cells (27, 70), while experiments in pig have shown that peripheral RPE cells respond to debridement of central RPE by proliferating (71). Indeed, preservation of the peripheral RPE is a prerequisite for successful RPE regeneration in the MRL/MpJ mouse model, which fails to regenerate RPE when high doses of sodium iodate cause degeneration of central and peripheral RPE (61, 72). Finally, the discovery of a subpopulation of RPE stem cells (30) suggests that an endogenous regeneration-capable population of RPE cells could exist in the human eye, and these might be analogous to injury responsive cells in zebrafish.

**Figure 9:**
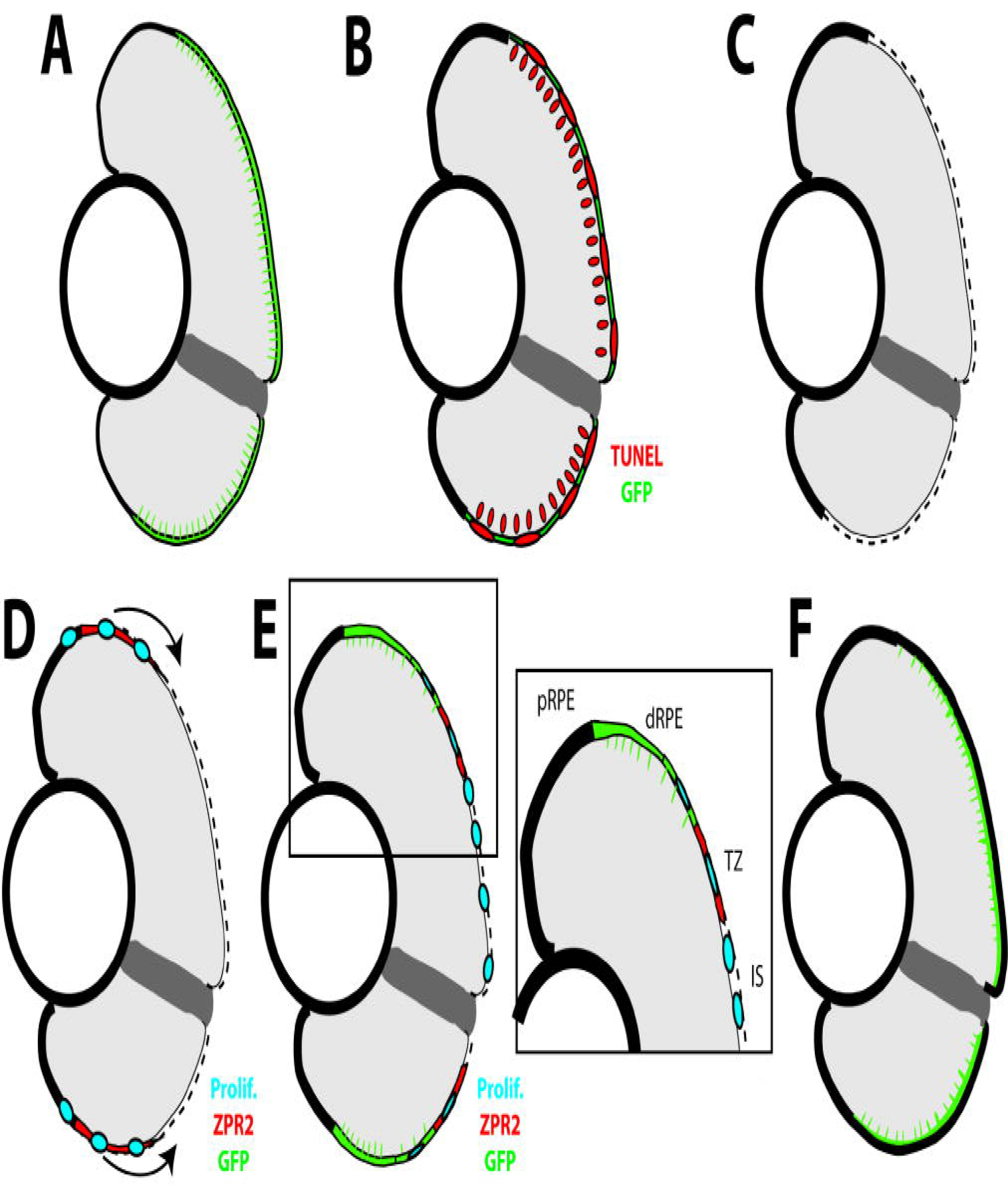
Model of RPE regeneration in larval zebrafish. (A) nfsB-GFP is specifically expressed in mature RPE in the central two-thirds of the eye. (B) Application of MTZ leads to apoptosis (red) of RPE and PRs. (C) RPE ablation leads to degeneration of PRs and Bruch’s Membrane (dotted line). (D) Unablated RPE in the periphery begin to proliferate and extend into the injury site (blue). (E) As regenerated GFP^+^ RPE appear in the periphery, the RPE can be divided into 4 zones: peripheral RPE, differentiated RPE, transition zone, and injury site. (E, inset) Regenerated differentiated RPE appears in the periphery proximal to the unablated peripheral RPE, and contains proliferative cells adjacent to the transition zone. The transition zone consists of still-differentiating RPE cells and proliferative cells. The injury site is comprised of unpigmented proliferative cells that do not express any RPE differentiation markers. (F) Regeneration of a functional RPE layer and Bruch’s Membrane is complete by 14dpi.

While we present strong evidence supporting a model in which regenerated RPE derives from injury-adjacent RPE cells, we cannot definitively establish the source of regenerated RPE without performing lineage tracing. In response to retinal injury, Muller glia proliferate and generate MGPCs that differentiate into new neurons (33). MGPCs are present at identical time points and in regions adjacent to ablated RPE. While it is possible that MGPCs transdifferentiate into RPE, this ability is not supported by any published studies. Another possible source for regenerated RPE is the CMZ, which generates neurons throughout the life of the animal (73–75). It was recently shown that *rx2*^+^ stem cells in the CMZ generate both RPE and retinal neurons (76). Thus, *rx2*^+^ cells in the CMZ could potentially respond to RPE injury by generating RPE in the periphery that migrate and proliferate within the injury site. Attempts at lineage tracing to date have been unsuccessful and therefore, it will be necessary to develop new genetic tools to unambiguously identify the source of regenerated RPE.

Mechanistically, we demonstate that the Wnt pathway is activated after RPE injury and that inhibiting Wnt impairs regeneration. While we have not identified the *lef1*-expressing cell type, two possibilities are: 1) apoptotic cells in the injury site and 2) cells of the immune system. There are notable similarities between *lef1* expression and TUNEL staining in the ONL and the RPE, and we detect *lef1* expression (Fig. 8B,B’) at the same time that TUNEL^+^ cells peak in ablated larvae (Fig. 1D-F). A previous study in *Hydra* showed that apoptotic cells are a source of Wnt3, and are required for head regeneration (77). The immune system has been shown to play a critical role in influencing the regenerative response (78) and Wnt signaling regulates the inflammatory properties of immune cells (79). Based on our results and previous data (80), we anticipate these responses to regeneration (e.g. apoptosis and infiltration of immune cells) to initiate immediately post-injury, which may explain why pre-treatment with IWR-1 does not affect regeneration. Wnt signaling has been shown to be important for RPE development *in vivo* (81, 82), thus it is also possible that Wnt activation is important at multiple times during RPE regeneration. Further work is required to determine how Wnt signaling modulates RPE regeneration.

Finally, we observed degeneration and regeneration of BM following ablation. That RPE ablation leads to BM breakdown may provide insight into the mechanisms underlying the initiation of CNV at the onset of exudative AMD. During CNV, choroidal endothelial cells penetrate BM and grow into the subretinal space; whether this process is initiated by the degeneration of the choriocapillaris or the RPE remains controversial (83, 84). Our results suggest that the RPE is required for BM maintenance. A logical next step would be to determine if choroidal vasculature invades the subretinal space following degeneration, as this would provide evidence that RPE degeneration is causative of CNV. Additionally, we show that zebrafish RPE are capable of repairing BM during regeneration. In human, BM undergoes a series of changes during aging that are thought to underlie AMD pathogenesis and inhibition of RPE function, changes that are thought to underlie the barrier to development of successful RPE transplant therapies (85, 86). The mechanisms underlying BM repair in zebrafish may provide critical insights into improving transplant survival and reintegration in humans.

## MATERIALS AND METHODS

### Fish maintenance and husbandry

Zebrafish were maintained following established protocols (see Supplementary Materials and Methods).

### RPE Ablation

Embryos derived from *rpe65a*:nfsB-GFP x AB crosses were maintained in PTU-containing (Sigma-Aldrich) fish water between 1-5dpf, and dechorionated at 3dpf. At 5dpf, larvae were exposed to 10mM MTZ (Sigma-Aldrich), dissolved in fish water, for 24 hours in the dark. After treatment, larvae were removed from MTZ and allowed to recover in fish water for 48 hours. At this point, severely ablated embryos were selected based upon the disruption of GFP signal in the RPE and lack RPE pigmentation. All ablated fish were then placed on a recirculating water system (Aquaneering) for recovery.

### Immunohistochemistry, TEM and *in situ* hybridization

Tissue preparation, immunohistochemistry, TEM, and *in situ* hybridization were performed by following established protocols (see Supplementary Materials and Methods).

### OKR Assays

Larvae were immobilized in 3% methylcellulose, oriented dorsal up and exposed to a full field rotating stimulus projected onto a screen (#NP100, NEC, Itasca, Illinois) that encompassed 180 degrees of the stimulated eye’s field of vision. Responses were captured using infrared light (880nm; #BL34-880, Spectrum, Montague, MI) through a Flea3 Camera (#FLU-U3-13Y3M-C, Point Grey Research, Richmond, BC, Canada) mounted on a dissecting microscope (#S8 APO, Leica Microsystems, Wetzlar, Germany). Videos were recorded by FlyCapture software (Point Grey) and quantified using custom MATLAB scripts (40).

### OCT imaging

Larvae were immobilized dorsal up in 3% methylcellulose and imaged with Optical Coherence Tomography (OCT) (∼840nm; Leica Bioptigen R2210 Spectral Domain Ophthalmic Imaging System). After imaging, larvae were rinsed and transferred into petri dishes to recover. OCT scans were analyzed using Bioptigen software (InVivoVue; Bioptigen, Research Triangle Park, NC), FIJI, and MetaMorph (Molecular Devices).

### Experimental design and statistics

To quantify transverse cryosections, three sections including or immediately adjacent to the optic nerve were quantified from each individual. Additional details on quantification and statistical analyses are included in in Supplementary Materials and Methods, and Supplemental Table 1. Data generated from immunohistochemical analyses are depicted in box and whisker plots, except for centrality measurements in Fig. 7K and S12C, in which all data are shown. In box and whisker plots, the box extends from the 25th to the 75th percentile and whiskers extend from the 5th to 95th percentile. Within the box, the median is indicated by a line and the mean is indicated by a “+”. Individual points outside the whiskers are plotted directly.

## ACKNOWLEDGEMENTS

We are grateful to Dick Barrett and Joshua Homme for technical assistance. This work was supported by a grant from the Macular Degeneration Research Program of the Bright Focus Foundation (M2016067), the Charles and Louella Snyder Retinal Regeneration Fund, and the E. Ronald Salvitti Chair in Ophthalmology Research to JMG, the Martha Wandrisco Neff Research Award in Macular Degeneration to NH and LLL, as well as NIH CORE Grant P30 EY08098 to the Department of Ophthalmology, the Eye and Ear Foundation of Pittsburgh, and from an unrestricted grant from Research to Prevent Blindness, New York, NY.

